# Region-specific features of early glial activation and Aquaporin-4 dysregulation in conditional mouse models of TDP-43 proteinopathies

**DOI:** 10.64898/2026.01.14.699494

**Authors:** Gabriela Nieva, Florencia Vassallu, Amaicha Depino, Vanina Netti, Lionel Muller Igaz

**Author notes:** Correspondence: Dr. Lionel Muller Igaz. Paraguay 2155 7mo piso, Buenos Aires, Argentina.

## Abstract

Aggregation and cytoplasmic mislocalization of TDP-43 are key features of several neurodegenerative diseases, including amyotrophic lateral sclerosis (ALS) and frontotemporal dementia (FTD). Neuroinflammatory processes mediated by glial cells play crucial roles in the pathophysiology of these and other diseases, defined as TDP-43 proteinopathies. Here, we characterized region-specific glial activation in two conditional mouse models: hTDP-43-WT (overexpressing nuclear wild-type human TDP-43) and hTDP-43-ΔNLS (expressing cytoplasmic TDP-43 with altered nuclear localization signal) following one month of transgene expression. Immunofluorescence analysis revealed distinct patterns of microglial activation across brain regions. hTDP-43-WT mice exhibited significant microgliosis in motor (MC) and somatosensory (SSC) cortices and hippocampal dentate gyrus (DG) with pronounced morphological alterations (i.e. increased soma size). Sholl analysis demonstrated reduced branching length and complexity in MC, SSC and hippocampal subfields. hTDP-43-ΔNLS mice displayed more pronounced microglial activation in hippocampal regions (CA1, DG) compared to cortical areas, with significant increases in microglial density. Additionally, we observed region-specific cortical astrocytosis in both models, suggesting coordinated glial reactivity. hTDP-43-ΔNLS mice showed decreased polarization of astrocytic water channel Aquaporin-4 (AQP4) around vascular structures in SSC and hippocampal CA1/DG. The changes in AQP4 localization, which is critical for glymphatic function, supports the hypothesis that this waste clearance system for the brain is altered in TDP-43 proteinopathies. These findings demonstrate that these different animal models of ALS/FTD induce distinct neuroinflammatory signatures, potentially contributing to the region-specific vulnerability observed in these diseases. Our data provide insights into early glial-mediated pathogenic mechanisms that could guide targeted therapeutic strategies for TDP-43 proteinopathies.

## Introduction

Neurodegenerative diseases are a heterogeneous group of progressive disorders characterized by selective neuronal vulnerability, accumulation of misfolded proteins, and irreversible functional decline. Despite their clinical diversity, conditions such as Alzheimer’s disease (AD), Parkinson’s disease (PD), amyotrophic lateral sclerosis (ALS), and frontotemporal dementia (FTD) share convergent pathogenic features, most notably the formation of disease-specific protein aggregates that disrupt cellular homeostasis and neuronal integrity (Kwon and Koh, 2020). In ALS and a substantial proportion of FTD cases, mislocalization and aggregation of TAR DNA-binding protein 43 (TDP-43) represent a defining molecular hallmark (Neumann et al., 2006), linking RNA dysregulation and proteostasis failure to neurodegeneration (Bright et al., 2021; McCauley and Baloh, 2019). Increasing evidence indicates that these proteinopathies are tightly coupled to innate immune activation within the central nervous system (CNS), where microglia and astrocytes undergo sustained phenotypic remodeling in response to chronic cellular stress (Haukedal and Freude, 2019). Both constitutive and conditional mouse models expressing wild-type or mutant TDP-43 variants have demonstrated that mislocalization of TDP-43 from the nucleus to the cytoplasm disrupts RNA metabolism, induces protein aggregation, and provokes widespread neuroinflammation (i.e. (Igaz et al., 2011), see review by (Armas et al., 2025)).

Neuroinflammation and reactive gliosis are now recognized as core components of neurodegenerative pathology across disease entities, detectable in patient biofluids, neuroimaging studies, post-mortem tissue, and experimental models (Bright et al., 2019; Kwon and Koh, 2020). Activated microglia and astrocytes can exert both neuroprotective and neurotoxic effects, influencing protein clearance, synaptic integrity, and neuronal survival in a context- and stage-dependent manner (Haukedal and Freude, 2019; Paolicelli et al., 2022).

Since the discovery of its involvement in neurodegenerative disease, the focus of research on TDP-43–driven pathology has been primarily on neuronal degeneration. However, more recent work increasingly shifted to other cellular populations and processes, with growing evidence pointing to neuroinflammation and non-neuronal cells as active, early contributors to pathology, not mere consequences (Provenzano et al., 2023). Glial cells, particularly microglia and astrocytes, are the primary orchestrators of this inflammatory response. Under pathological conditions, these cells undergo a transformation from homeostatic supporters to active participants in the disease process. However, their roles are complex and their activation phenotypes are highly context-dependent (Ajoolabady et al., 2025; Santiago-Balmaseda et al., 2024).

While the role of toxic astrocytes is well-established in SOD1-based animal models of ALS (Monteiro Neto et al., 2025; Provenzano et al., 2023), less is known about how TDP-43 pathology, which is relevant to most ALS cases, drives specific glial activation signatures. Pathological TDP-43 has been shown to engage inflammatory signaling pathways, including NF-κB, inflammasome activation, and cGAS–STING signaling (Bright et al., 2021; McCauley and Baloh, 2019; Swarup et al., 2011), while genetic risk factors for ALS/FTD converge on immune regulatory pathways (McCauley and Baloh, 2019), supporting a bidirectional and self-reinforcing relationship between protein aggregation and neuroinflammation. Together, these observations identify gliosis not merely as a secondary response to neuronal loss but as an integral driver of disease progression and a potential therapeutic target in TDP-43–associated neurodegeneration (Bright et al., 2021).

To elucidate the role of TDP-43 in triggering glial activation, we used two conditional transgenic mouse models with different patterns of TDP-43 expression in forebrain neurons: one overexpressing wild-type human TDP-43 with preserved nuclear localization (tTA/WT), and a second one expressing a human TDP-43 variant lacking the nuclear localization signal, leading to pathological cytoplasmic localization (tTA/ΔNLS) (Igaz et al., 2011). We hypothesized that these two TDP-43 species would induce different and region-specific early microglial and astroglial activation signatures, which might represent an early marker of disease progression. The one month post-transgene induction time window was selected to capture sustained glial changes prior to overt degeneration in these inducible TDP-43 models. Moreover, as ΔNLS mouse models have been shown to present more varied and severe phenotypes than TDP-43 WT expressing ones (Alfieri et al., 2014; Alfieri et al., 2016; Igaz et al., 2011; Walker et al., 2015), we also analyzed if Aquaporin-4 (AQP4) was dysregulated in the brain after neuronal cytoplasmic TDP-43 expression.

AQP4, the most abundant water channel in the brain, is an integral component of the glymphatic system. Its distribution is strongly polarized in the endfeet of astrocytes which ensheathe the blood vessels in the brain (Mader and Brimberg, 2019). This specific localization allows AQP4 to be in close contact with the perivascular space, thus facilitating CSF influx into the brain parenchyma, which is crucial for the effective functioning of the glymphatic system (Szlufik et al., 2024). In recent years, many studies have reported the pathophysiological role that AQP4 plays in several CNS neurodegenerative disorders such as PD and AD by affecting the clearance function of the glymphatic system (Peng et al., 2023; Szlufik et al., 2024) and the formation of protein aggregates such as α-synuclein (α-Syn) in PD and β-amyloid (Aβ) in AD, thus contributing to the pathogenesis of neurodegeneration. The loss of AQP4 perivascular localization (and the impairment of glymphatic flow) was reported in AD patients as well as in animal models (Szlufik et al., 2024). Here, we assessed the status of AQP4 polarization in the brains of tTA/ΔNLS mice to evaluate a potential contribution of glymphatic dysfunction to phenotype development in this model of TDP-43 proteinopathies.

Our results demonstrate that one month of wild-type hTDP-43 overexpression or cytoplasmic hTDP-43 expression is sufficient to induce significant cortical and hippocampal gliosis. Collectively, these findings emphasize the relevance of studying the early phases of the pathological processes associated with neurodegenerative conditions, and suggest that modulating these early events might eventually contribute to the development of therapeutic approaches for TDP-43 proteinopathies.

## Materials and methods

### Animals

All animal experiments were approved by the National Animal Care and Use Committee (CICUAL) of the School of Medicine, University of Buenos Aires (UBA). All efforts were made to minimize animal suffering and to reduce the number of animals used. Conditional transgenic mice expressing either wild-type human TDP-43 (hTDP-43-WT) or TDP-43 with a mutated nuclear localization signal (hTDP-43-ΔNLS) were generated as previously described (see (Alfieri et al., 2014; Alfieri et al., 2016; Igaz et al., 2011; Silva et al., 2019)). Both lines were backcrossed on a C57BL/6J background for at least 15 generations. Experimental groups consisted of animals positive for both the CaMKIIα-tTA and tetO-hTDP-43 transgenes (either WT or ΔNLS TDP-43 variants; from now on, tTA/WT and tTA/ΔNLS, respectively). The control group included non-transgenic littermates or those carrying only one of the transgenes (CaMKIIα-tTA or tetO-hTDP-43). Mice from both sexes were used, as in our previous studies (Alfieri et al., 2014; Alfieri et al., 2016; Igaz et al., 2011; Silva et al., 2019). Mice were housed in a pathogen-free facility under controlled environmental conditions (23 ± 2 °C; 40–60% relative humidity) with a 12 h light/dark cycle and ad libitum access to food and water. Environmental enrichment consisted of cardboard rolls and paper, replaced only when damaged or heavily soiled.

### Doxycycline treatment and transgene induction

To prevent prenatal and postnatal transgene expression, all breeding pairs and offspring received drinking water containing doxycycline hyclate (DOX, 0.2 mg/ml; sc-204734A, Santa Cruz Biotechnology, USA) from conception until weaning at postnatal day 28 (P28), as previously described (Alfieri et al., 2014). At P28, mice were switched to regular drinking water to induce transgene expression. Mice were analyzed at 2 months of age, corresponding to 1 month of continuous transgene induction.

### Genotyping

Ear biopsies were collected between postnatal days 15 and 20. DNA extraction and PCR amplification were performed in our Molecular Biology Laboratory using previously described primer sequences (Alfieri et al., 2014; Alfieri et al., 2016; Silva et al., 2019).

### Perfusion and tissue processing

Mice were deeply anesthetized with 5% chloral hydrate (1 ml / 30 g) and transcardially perfused with phosphate-buffered saline (PBS, 0.01 M, pH 7.4) supplemented with 10U/ml Heparin. The brains were immediately extracted and post-fixed overnight in 4% paraformaldehyde (PFA) at 4 °C, then cryoprotected in 10% and 30% sucrose in PBS. Coronal sections (50 µm) were obtained using a microtome (SM 2010R; Leica) and stored at −20°C in cryoprotecting solution (50% glycerol, 50% PBS).

### Immunofluorescence staining

Double immunofluorescence was performed as follows: free-floating sections were rinsed in PBS (0.01 M) and permeabilized for 1 h at room temperature (RT) in PBS containing 1% Triton X-100. Sections were then blocked for 1 h in PBS containing 5% normal goat serum (NGS) and 0.3% Triton X-100. The primary antibodies (diluted in 0.3% Triton X-100 and 3% NGS in PBS) were incubated overnight at 4°C with the indicated dilutions: rabbit anti-Iba1 antibody (1:3000; Wako) for microglial labeling, rabbit anti-GFAP antibody (1:2000; DAKO) for astrocytic labeling, and monoclonal mouse anti-hTDP-43 antibody (1:10000; Proteintech). The next day, sections were washed 3×5 min in PBS containing 0.04% Triton X-100 and incubated for 4 h at RT in the dark with fluorophore-conjugated secondary antibodies diluted in 3% NGS, 0.3% Triton X-100 in PBS: rhodamine-conjugated goat anti-rabbit IgG (1:500; Jackson Laboratories) and Alexa Fluor 488–conjugated goat anti-mouse IgG (1:500; Invitrogen). After three washes in PBS, nuclei were counterstained for 30 min at RT in the dark with Hoechst 33342 (1 µg/ml; Sigma-Aldrich) and washed twice in PBS. Sections were mounted on glass slides and coverslipped with 30% glycerol in PBS. Slides were stored at 4 °C in the dark until imaging.

For AQP4/GFAP double immunofluorescence staining, brain tissue slices were permeabilized with 0.3% Triton X-100 in PBS for 15 minutes at RT and blocked with BSA 1mg/ml (Sigma-Aldrich) in PBS with 0.03% Triton-X-100 for 1h at RT. Samples were incubated with a rabbit anti-AQP4 primary antibody (1:/300; sc-20812 Santa Cruz) overnight at 4°C, followed by three washes in 0.1% Triton X-100/PBS and then incubated with secondary Alexa 488 donkey anti-rabbit antibody (1:300; 711-545-152, Jackson Laboratories) for 2 hours at RT. After washes, slices were incubated with a rat anti-GFAP antibody (1:500, 2.2B10, generously provided by Dr. Virginia Lee, University of Pennsylvania) overnight at 4°C and then with a secondary Alexa 647 goat anti-rat antibody (1:1000, Invitrogen) for 2 hours at RT. Finally, cell nuclei were identified using Hoechst 33342 (1 µg/ml; Sigma-Aldrich) for 30 minutes at RT. Coverslips were mounted with Vectashield mounting medium (VectorLabs).

### Image acquisition and analysis

Fluorescent images were acquired using a Zeiss AxioImager 2 microscope equipped with the structured illumination system ApoTome 2 (Carl Zeiss, Germany), using a Hamamatsu Orca Flash 4.0 camera, under identical settings for all groups. For experiments with both tTA/WT and tTA/ΔNLS mice, two separate immunofluorescences were performed: one for Iba1 and one for GFAP, sometimes combined with hTDP-43 staining. The number of animals analyzed varied depending on the immunostaining and the brain region examined and are indicated in the corresponding figure legends.

Microglial analysis from Iba1-stained sections included Iba1-positive (Iba+) area (normalized to the mean of the control group), microglial density (Iba1+ somas/mm²), and soma area of Iba1+ cells (µm²). Sholl analysis parameters included the number of intersections as a function of distance from the soma (µm), reflecting branching complexity; the total number of intersections; the total process length (µm), representing overall arbor size; and the maximum radial process extension (µm). Astrocytic reactivity in GFAP-stained sections was additionally quantified as GFAP-positive area (normalized to the mean of the control group).

All measurements, image processing and quantification were performed using Fiji (Schindelin et al., 2012). Images for Iba1 and GFAP quantifications were captured at 20X magnification (N.A. 0.8 objective). For Sholl analyses, z-stacks of Iba1-stained microglia were acquired at 40X (N.A. 0.95 objective) using 1 µm step size, typically yielding 30–45 optical sections per sample. Maximum-intensity projections were generated in Fiji prior to binarization, skeletonization, and subsequent morphometric analysis.

AQP4 polarization was quantified as previously described (Hablitz et al., 2020; Wu et al., 2025) using 40X images acquired as described for GFAP and Iba1 staining. Briefly, 50 μm segments centered on blood vessels (as identified by vascular-shaped AQP4 localization) were analyzed using the line-plot tool in ImageJ. 5 vessels (range of 5–8 μm wide) were chosen per image (2 per region in each animal). The baseline fluorescence was defined as the average intensity over 10 μm, −20 μm to −10 μm from peak fluorescence. AQP4 polarization index (PI) was calculated as the ratio between the mean vascular endfeet AQP4 fluorescent signal and mean background signal. Average polarity was calculated and the PI of each group was normalized to the Control group. The higher the AQP4 PI, the greater proportion on immunoreactivity was restricted to perivascular regions, whereas the lower PI, the more evenly distributed immunoreactivity was between the perivascular endfeet and the cell body. Blood vessel diameter was determined from line-plots by measuring the distance between the initial and terminal high-intensity AQP4 fluorescence signals, corresponding to the start and end of each vessel.

### Statistical analysis

Graph Pad Prism 8 software (GraphPad Software, Inc., San Diego, CA, USA) was used for data visualization and statistical analysis. Comparisons between two groups for each brain region were made using unpaired two-tailed Student’s *t*-test. Sholl data were analyzed by two-way ANOVA with repeated measures followed by Sidak’s multiple comparisons test. Data are expressed as mean ± standard error mean (SEM), and significance was set at *p* < 0.05.

## Results

### Wild-type hTDP-43 overexpression induces widespread microglial activation and morphological remodeling

Although microglial activation is a feature of several TDP-43 murine models of proteinopathies (Igaz et al., 2011; Jara et al., 2019), the specific regional contribution in early post induction stages has not been investigated. We observed a significant increase in microglial activation across multiple cortical and hippocampal regions after one month of transgene expression in tTA/WT hTDP-43 mice (Figure 1). Immunofluorescence analysis confirmed robust neuronal expression (Igaz et al., 2011; Silva et al., 2019) of human wild-type TDP-43 with predominant nuclear localization across cortical and hippocampal regions in tTA/WT mice, while no signal was detected in control littermates (Figure 1A), confirming neuronal transgene expression and validating lack of colocalization with microglia, as expected. Quantitative analysis of immunofluorescence stained sections revealed a significant elevation in the total immunopositive area for Iba1 in the motor cortex (MC, +21.0%), somatosensory cortex (SSC, +21.0%), and dentate gyrus (DG, +28.0%) compared to control littermates (Figure 1B). This was accompanied by a significant increase in microglial density (Iba1+ somas/mm²) in the same regions (MC: +19.0%, SSC: +33.2%, DG: +33.5%) (Figure 1C). Notably, when the CA1 region was analyzed as a whole, no significant differences in Iba1-immunopositive area were detected between groups. However, given the well-defined laminar organization of the hippocampal CA1, we performed a layer-specific analysis (Supplementary Figure 1). This approach revealed a significant increase in Iba1+ area in the stratum oriens (Or) and stratum radiatum (Rad) of tTA/WT hTDP-43 mice, whereas no changes were observed in the pyramidal layer (Py) (Supplementary Figure 1C). DG subregional analysis revealed significant increases for both the granular (Gr) and the polymorphic (Po) layers (Supplementary Figure 2D). A key indicator of microglial activation, soma hypertrophy, was evident through a significant enlargement of Iba1+ soma area in the prefrontal cortex (PFC: +15.2%), MC (+13.7%), SSC (+12.5%), and DG (+13.8%) (Figure 1D).

**Figure 1.**
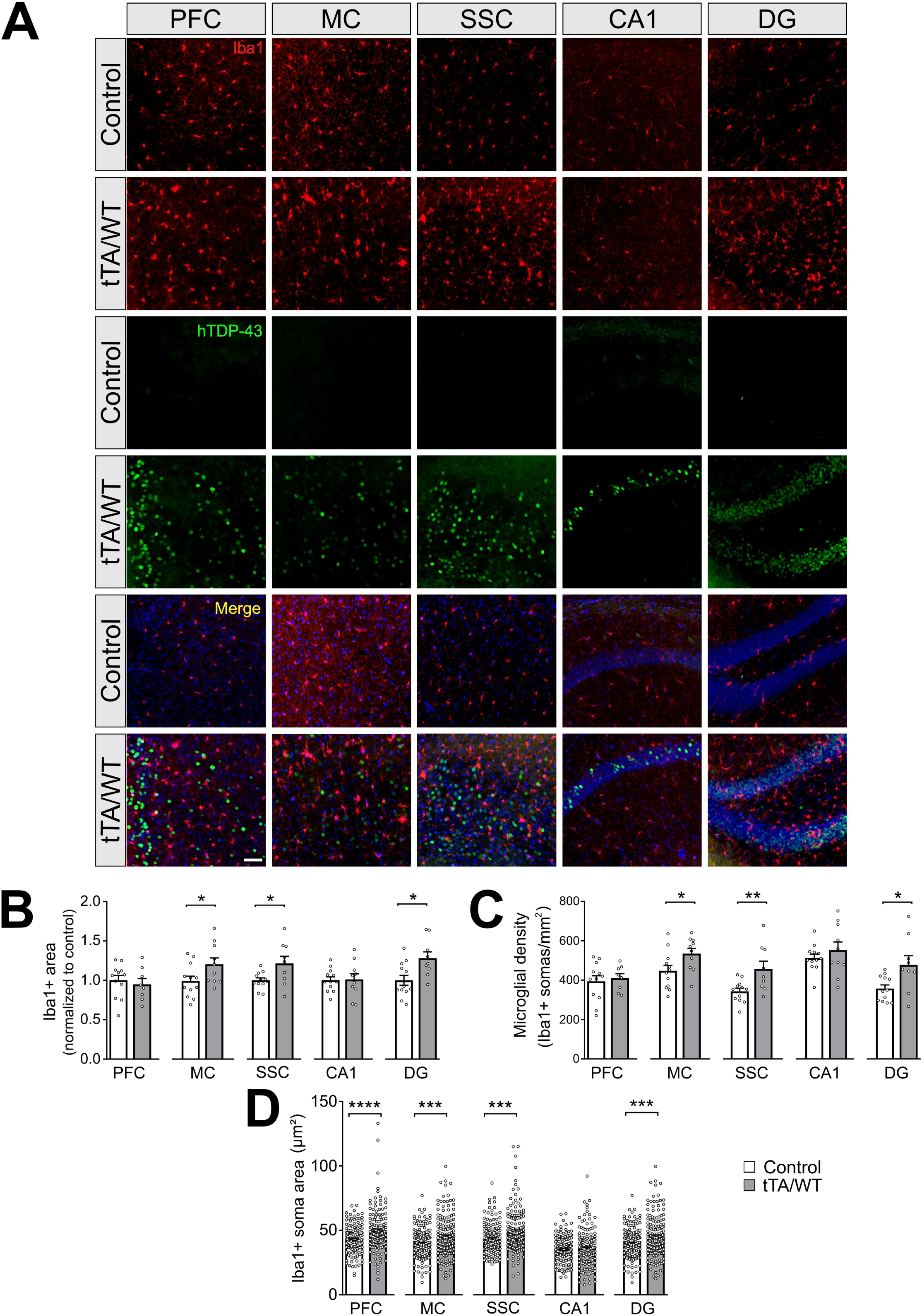
Widespread microglial activation across cortical and hippocampal regions following transgene expression in tTA/WT hTDP-43 mice. **(A)** Representative immunofluorescence images show Iba1+ microglia (red), human TDP-43 (green), and merged channels (plus nuclear marker Hoechst in blue) in the prefrontal cortex (PFC), motor cortex (MC), somatosensory cortex (SSC), CA1 hippocampal region, and dentate gyrus (DG) of control and tTA/WT mice after one month of transgene expression. Scale bar: 50 µm. **(B-D)** Quantitative analyses reveal a significant increase in Iba1+ area (normalized to control) **(B)** and microglial density (Iba1+ somas/mm²) **(C)** in the MC, SSC, and DG of tTA/WT mice compared with controls. Moreover, Iba1+ microglia exhibit enlarged soma area (µm²) **(D)** in the PFC, MC, SSC, and DG, a hallmark of microglial activation. Statistical significance was determined by Student’s t test (* p < 0.05; ** p < 0.01; *** p < 0.001; and **** p < 0.0001). Results are expressed as mean ± SEM. n = 8 - 12 animals per group (B-C); n = 144 - 175 cells per group (D) from the same number of animals as in B-C.

In order to further evaluate morphological changes beyond cell size and density, we performed Sholl analysis on reconstructed microglial cells (Figure 2). In tTA/WT hTDP-43 mice, microglia exhibited a significant reduction in process complexity. This was evidenced by a lower number of process intersections at increasing distances from the soma in the MC, SSC, CA1, and DG hippocampal regions (Figure 2C-G, panel i). Consistent with this, we found a decrease in the total number of intersections (MC: -26.1%, SSC: -26.3%, CA1: -46.2% and DG: -39.3%) and total process length (MC: -28.1%; SSC: -25.5%, CA1:-48.2% and DG: -42.9%) in these areas (Figure 2C-G, panels ii and iii). Maximum process extension was significantly lower in MC (-21.1%) and hippocampal CA1 (-35.9%) and DG (-41.1%) regions (Figure 2C-G, panel iv). This shift towards a less ramified, more hypertrophied or bushy morphology, characterized by process retraction, is a well-established feature of microglial activation (Green and Rowe, 2024).

**Figure 2.**
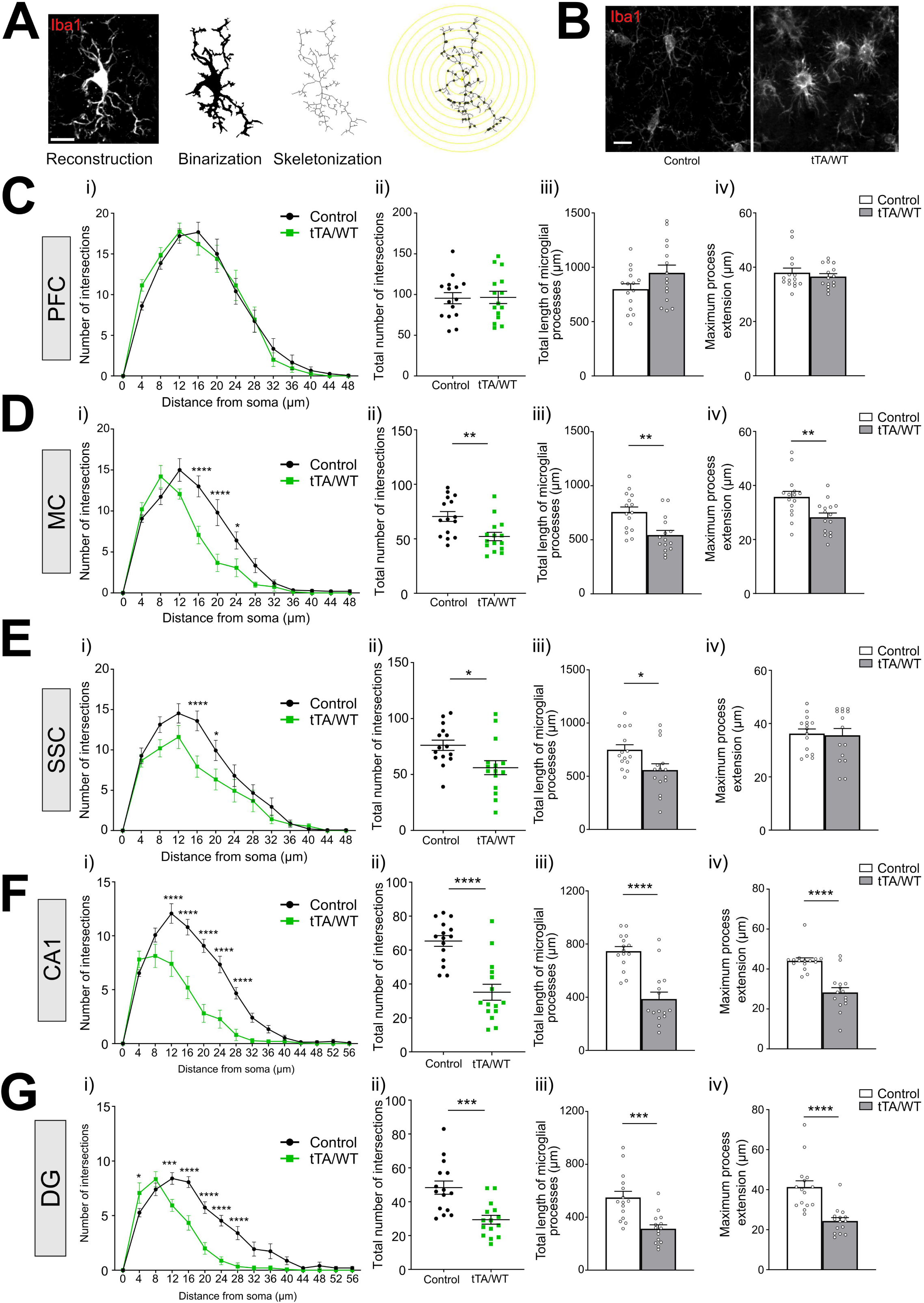
Sholl analysis reveals reduced microglial process complexity in multiple brain regions of tTA/WT hTDP-43 mice. **(A)** Z-stacks were acquired using a Zeiss AxioImager 2 ApoTome 2 system with a 40× objective and processed as maximum intensity projections. For each animal, five Iba1+ microglial cells were randomly selected, reconstructed in Fiji (ImageJ, NIH, USA), binarized, and skeletonized for process tracing. Sholl analysis quantified intersections of microglial processes with concentric circles radiating from the soma. Scale bar = 10 µm. **(B)** Representative maximum intensity Z-stack images of microglia from the MC regions of control and tTA/WT mice. Scale bar = 10 µm. **(C-G)** Quantitative Sholl analysis of Iba1+ microglia in the prefrontal cortex (PFC), motor cortex (MC), somatosensory cortex (SSC), CA1 hippocampal region (CA1), and dentate gyrus (DG) demonstrate significant morphological alterations after one month of transgene expression in tTA/WT mice compared with control littermates. Sholl profiles indicated a reduction in the number of intersections at increasing distances from the microglial soma **(i)**, total number of intersections **(ii)** and total process length **(iii)** in MC, SSC, CA1 and DG hippocampal regions. Maximal process extension **(iv)** was reduced in tTA/WT mice, consistent with microglial activation and neuroinflammation. Statistical analysis of Sholl profiles (number of intersections vs. distance) was performed using two-way ANOVA followed by Sidak’s multiple comparisons test (* p < 0.05; ** p < 0.01; *** p < 0.001), whereas total number of intersections, total process length, and maximal process extension were analyzed using Student’s t-test (* p < 0.05; ** p < 0.01; *** p < 0.001; **** p < 0.0001); n = 15 cells/3 mice per group analyzed. Results are expressed as mean ± SEM.

### Cytoplasmic hTDP-43 expression elicits a more pronounced microglial activation phenotype

We next investigated whether mice expressing the cytoplasmic TDP-43 variant would elicit a different microglial response. In tTA/ΔNLS hTDP-43 mice, we observed robust microglial activation and soma hypertrophy across several cortical and hippocampal areas analyzed (Figure 3). Specifically, there were significant increases in microglial density in the MC (+30.9%), CA1 (+73.8%), and DG (+157.4%) regions (Figure 3C) and immunopositive area for Iba1 in the same areas plus the SSC (MC: +46.9%, SSC: +25.6%, CA1: +61.7%, DG: +53.1%) (Figure 3B) compared to controls. Of note, microglial soma size was significantly increased in all three cortical areas but not in hippocampal subregions (PFC: +102.5%, MC: +23.9%, SSC: +9.8%) (Figure 3D).

**Figure 3.**
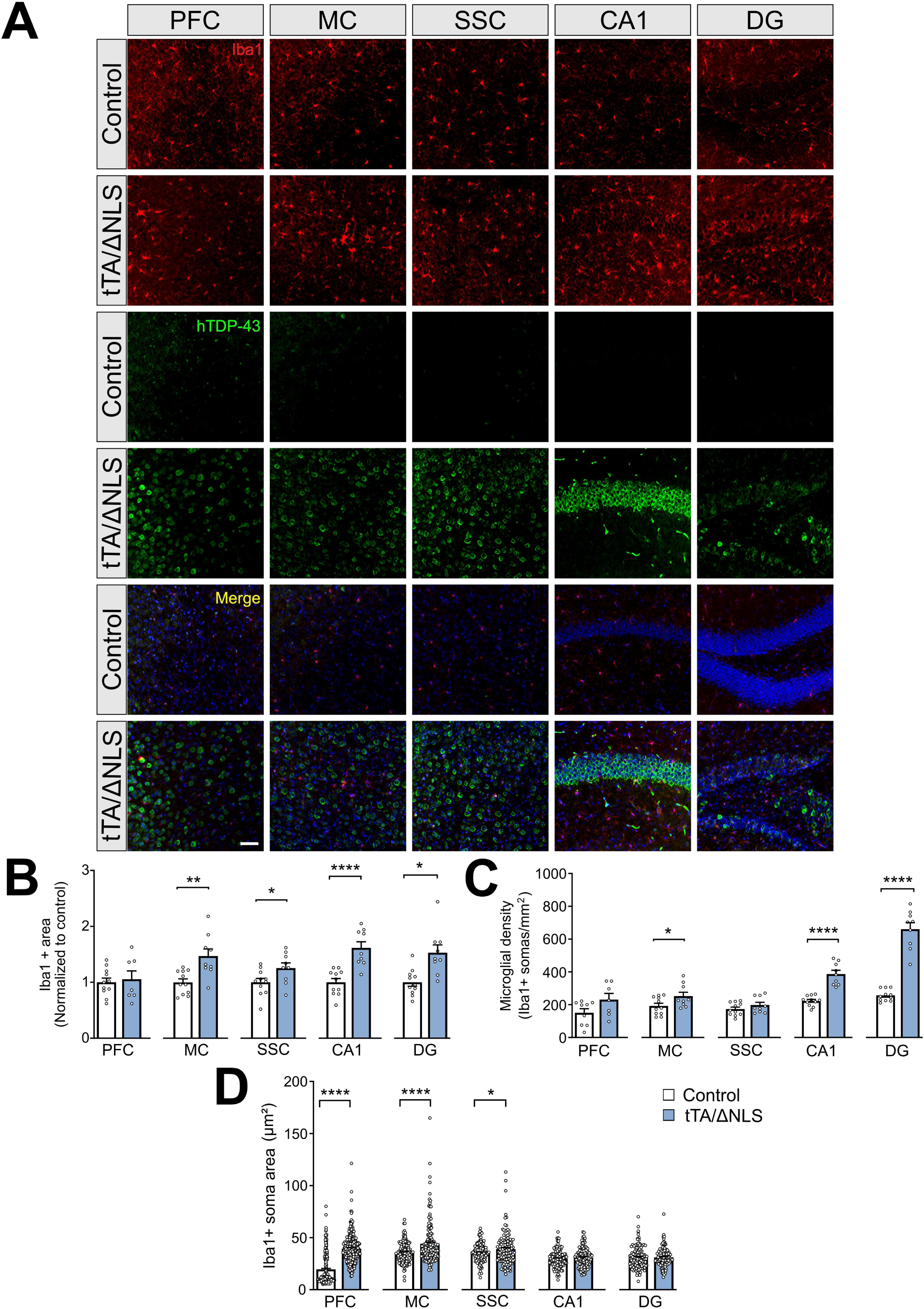
Microglial activation and soma hypertrophy across cortical and hippocampal regions in tTA/ΔNLS hTDP-43 mice. **(A)** Representative immunofluorescence images show Iba1+ microglia (red), human TDP-43 staining (green), and merged channels (plus nuclear marker Hoechst in blue) in the prefrontal cortex (PFC), motor cortex (MC), somatosensory cortex (SSC), CA1 hippocampal region, and dentate gyrus (DG) of control and tTA/ΔNLS mice after one month of transgene expression. Scale bar: 50 μm. Significant increases in Iba1+ area **(B)** and microglial density **(C)** are observed in the MC, CA1, and DG regions in the tTA/ΔNLS hTDP-43 animals compared to control animals after one month of transgene expression. Iba1+ microglia also demonstrate significantly increased soma size **(D)** in the PFC, MC, SSC regions, which is a key morphological hallmark of neuroinflammation. The analysis was based on images acquired using a Zeiss AxioImager 2 ApoTome 2 system at a 20X objective. Statistical significance was determined by Student’s t test (* p < 0.05; ** p < 0.01; **** p < 0.0001). Error bars represent SEM. n = 7 - 11 animals per group (B-C); n = 122 - 263 cells per group (D) from the same number of animals as in B-C.

To resolve whether microglial activation was uniformly distributed within hippocampal subfields, we conducted a layered analysis of Iba1 immunoreactivity to assess whether the increased Iba1 immunoreactivity observed in tTA/ΔNLS mice reflected uniform microglial changes across hippocampal layers (Supplementary Figure 2). Layer-specific analysis revealed a significant increase in Iba1-positive area in CA1 strata and in the granule cell layer of the DG, whereas no significant differences were detected in the polymorphic layer (Supplementary Figure 2C-D). These results indicate that microglial activation induced by cytoplasmic hTDP-43 is spatially different at the laminar level. This selectivity supports the notion of distinct neuroinflammatory signatures driven by nuclear and cytoplasmic TDP-43 overexpression.

Sholl analysis in the tTA/ΔNLS hTDP-43 animals revealed an extensive and profound decrease in microglial ramification, indicating a transition to an activated, amoeboid-like state (Figure 4B-F, panel i). Across MC, SSC, CA1, and DG, microglia showed a significant reduction in process complexity, total number of intersections (MC: -49.9%, SSC: -54.9%, CA1: -46.1%, and DG: -78.0%), and total process length compared to controls (MC: -52.9%, SSC: -55.0%, CA1: -44.8%, and DG: -77.1%) (Figure 4B-F, panels ii, iii and iv). This generalized process retraction suggests a more severe and widespread activated phenotype compared to that induced by the expression of WT-hTDP-43 transgene.

**Figure 4.**
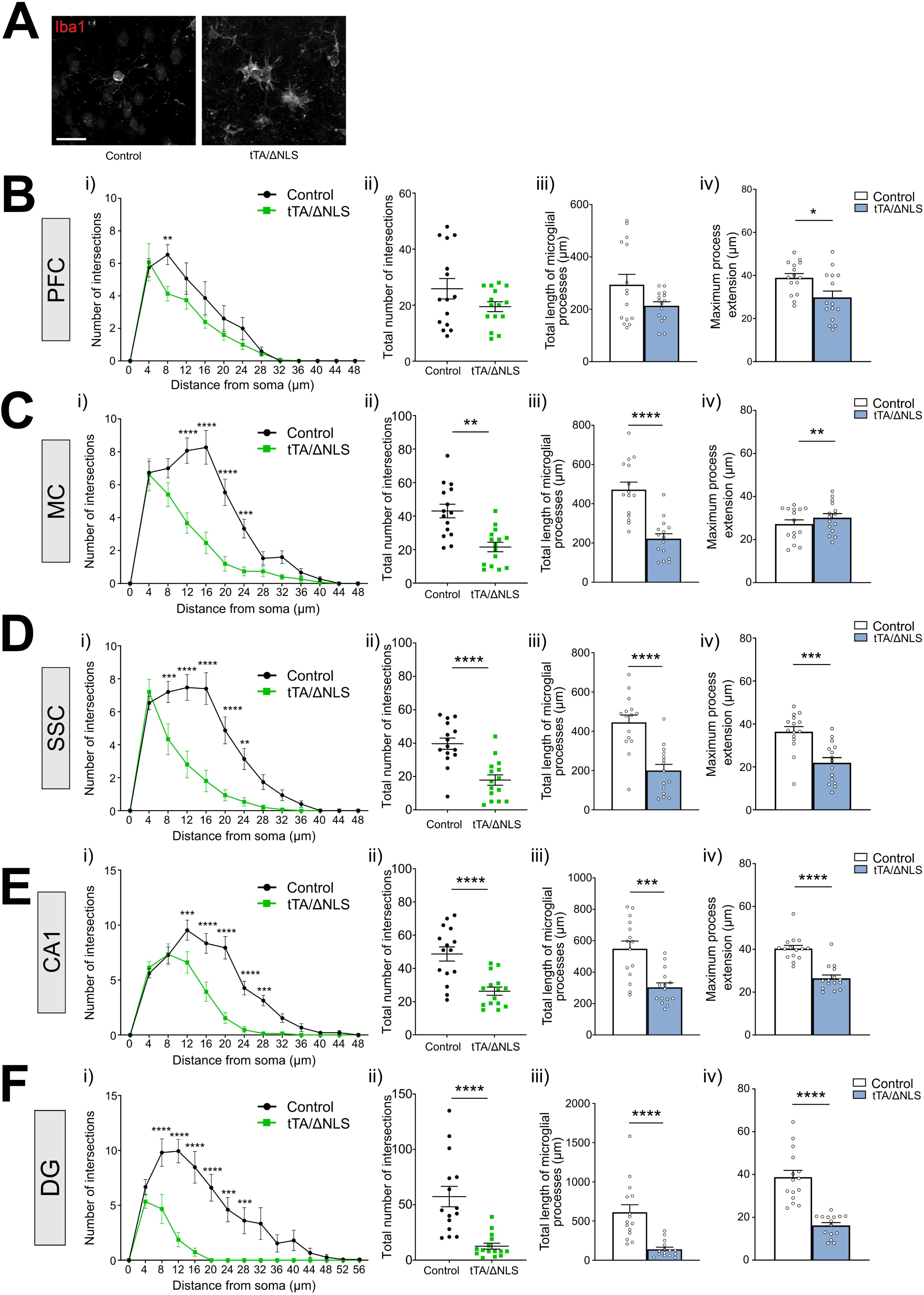
Microglial ramification is significantly impaired across multiple brain regions in tTA/ΔNLS hTDP-43 mice after one month of gene expression, indicating an activated/amoeboid state. **(A)** Representative maximum intensity Z-stack images of microglia from the MC of control and tTA/ ΔNLS mice. Scale bar = 20 µm. **(A-F)** Sholl analysis performed in prefrontal cortex (PFC), motor cortex (MC), somatosensory cortex (SSC), CA1 hippocampal region (CA1), and dentate gyrus (DG) demonstrates a significant reduction in microglial process complexity **(i)**, total number of intersections **(ii)**, and total length of microglial processes **(iii)** in the tTA/ΔNLS hTDP-43 animals compared to control littermates. This reduced ramification (microglial process retraction) **(iv)** is a key morphological hallmark of neuroinflammation and transformation to an activated/amoeboid phenotype. The analysis was performed after one month of transgene expression. Images used for quantification were acquired using a Zeiss AxioImager 2 ApoTome 2 system with a 40X objective. Statistical analysis of Sholl profiles (number of intersections vs. distance) was performed using two-way ANOVA followed by Sidak’s multiple comparisons test (** p < 0.01; *** p < 0.001, *** p < 0.0001), whereas total number of intersections, total process length, and maximal process extension were analyzed using Student’s *t*-test (* p < 0.05; ** p < 0.01; *** p < 0.001; **** p < 0.0001);. n = 15 cells/3 mice per group analyzed. Results are expressed as mean ± SEM.

### Differential astroglial activation highlights a widespread response in tTA/ΔNLS hTDP-43 mice

Although reactive astrocytosis is a key hallmark of many neurodegenerative diseases (Brandebura et al., 2023), most studies in ALS/FTD mouse models focus on the glial activation in the spinal cord. To determine the early response of the astrocyte population in the brains of our inducible TDP-43 mouse models, we quantified the area covered by GFAP+ cells (Figure 5). One month after induction, GFAP-based astrogliosis patterns differ between WT and ΔNLS models. Astroglial activation was selective and more regionally restricted in hTDP-43-WT mice, with GFAP upregulation confined primarily to MC (+363.0%) and SSC (+199.0%) (Figure 5B). In contrast, tTA/ΔNLS mice displayed a significant and widespread astroglial activation across all cortical and hippocampal regions analyzed (PFC: +92.58%, MC: +259.0%, SSC: +215.5%, CA1: +50.4%, and DG: +51.3%) (Figure 5C). This robust astroglial reactivity indicates that the presence of cytoplasmic TDP-43 triggers a more pervasive neuroinflammatory response from astrocytes compared to the wild-type protein.

**Figure 5.**
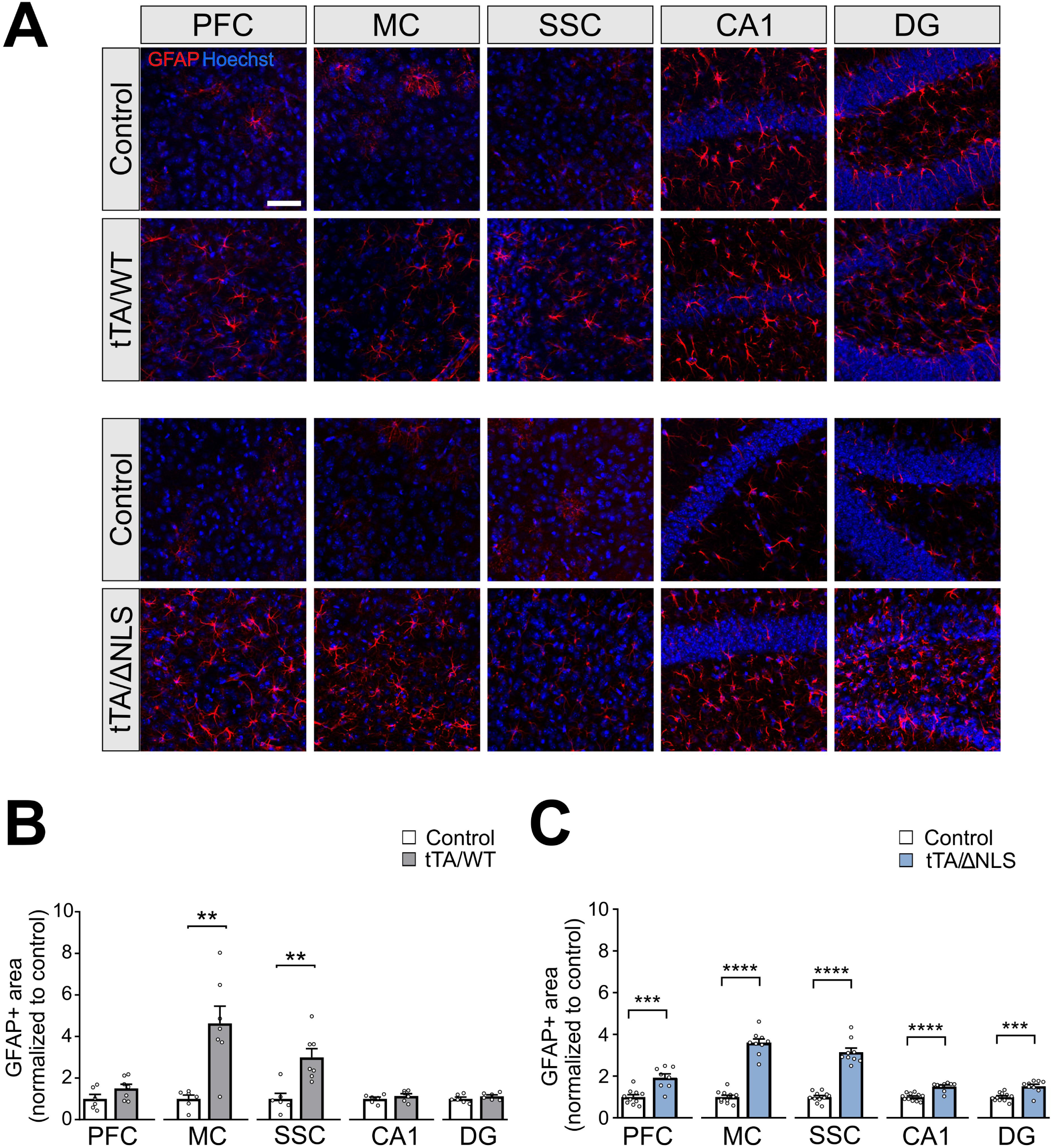
Region-specific astroglial activation in tTA/WT and widespread activation in tTA/ΔNLS hTDP-43 mice. **(A)** Representative immunofluorescence images show GFAP+ astrocytes (red) and nuclear marker Hoechst (blue) in control and tTA/WT mice (top two rows), and in control and tTA/ΔNLS hTDP-43 mice (two bottom rows), across the prefrontal cortex (PFC), motor cortex (MC), somatosensory cortex (SSC), CA1 hippocampal region (CA1), and dentate gyrus (DG). Scale bar: 50 µm. **(B-C)** Quantification of GFAP+ area (normalized to control) revealed selective astroglial activation in MC and SSC of tTA/WT mice **(B)**, whereas tTA/ΔNLS hTDP-43 animals exhibited significant increases in GFAP+ area across all analyzed regions **(C)**, consistent with robust astroglial reactivity and neuroinflammation. Images were acquired using a Zeiss AxioImager 2 ApoTome 2 system. Statistical analysis was performed using Student’s *t*-test (** p < 0.01; *** p < 0.001; **** p < 0.0001). Results are expressed as mean ± SEM. n = 6 - 7 animals per group (B); n = 8 - 11 animals per group (C).

### Decreased AQP4 Polarization Index (PI) in SCC and hippocampus of hTDP-43-ΔNLS animals

To investigate whether glymphatic system function, which relies on astrocytic AQP4 expression, is altered in hTDP-43-ΔNLS mice, we evaluated AQP4 expression and polarization, as well as the diameter of blood vessels in selected brain regions (SSC, CA1 and DG). We focused on this animal model given its more widespread astroglial activation, which provided an appropriate context to explore early alterations in astrocytic AQP4 polarization. Figure 6 shows AQP4 (green channel) and GFAP (red channel) expression and the merged channel. AQP4 is highly polarized to perivascular astrocytic end feet in control animals, but the AQP4 PI was reduced in SSC (-18.5%), CA1 (-32.6%) and DG (-46.0%) of hTDP-43-ΔNLS mice. Blood vessel diameter was reduced only in DG of these animals, likely due to higher rate of neurodegeneration reported in this area (Igaz et al., 2011). No changes in AQP4 total intensity were observed in CA1, DG or SSC between control or hTDP-43-ΔNLS mice (not shown). This early decrease in AQP4 polarization suggests that mice expressing cytoplasmic TDP-43 likely present an early impairment in perivascular astrocytic organization.

**Figure 6.**
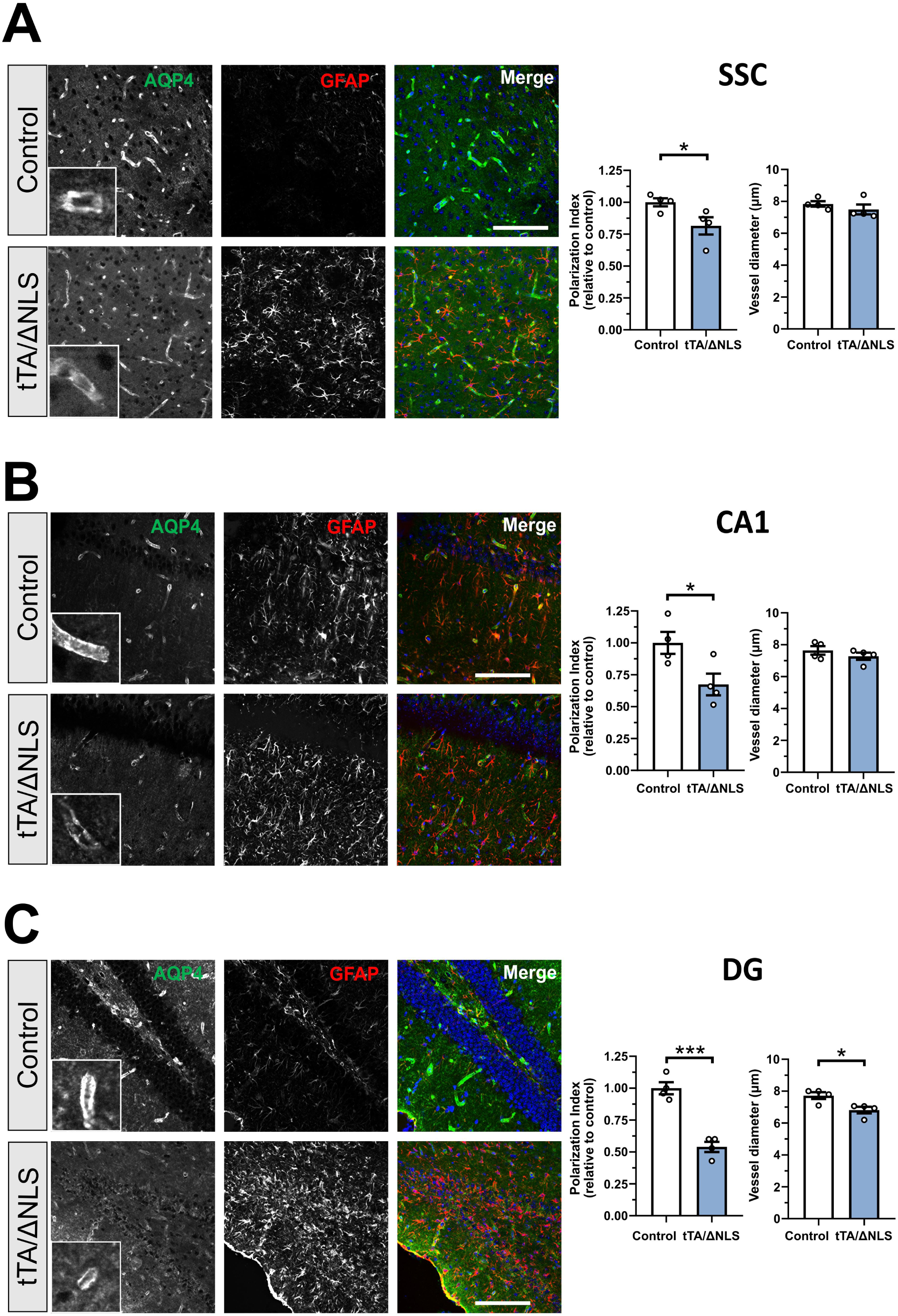
Perivascular AQP4 polarization is decreased in the cortex and hippocampus of tTA/ΔNLS hTDP-43 mice. hTDP-43-ΔNLS mice show decreased AQP4 polarization index (PI) around vascular structures in SSC and hippocampal CA1/ DG in comparison to control animals. Representative images of AQP4 (green channel), GFAP (red channel) and merged channel (plus nuclear marker Hoechst in blue) in SSC **(A)**, CA1 **(B)** and DG **(C)**. Images are representative of 4 animals (10 vessels/animal) in 2 independent experiments (Scale bar: 100 μm). Inserts show enlarged boxed areas from the green channel image (inset : 28 μm). Quantification of AQP4 polarization index (PI) and vessel diameter (scale bar: 28 μm). PI was calculated as the ratio between the mean vascular end foot AQP4 fluorescent signal and mean background signal. Results are expressed as mean ± SEM. * p < 0.05, *** p < 0.0001 control vs. hTDP-43-ΔNLS mice (Student’s test).

## Discussion

In this current study we provide a detailed characterization of early, region-specific glial responses to neuron-specific expression of different TDP-43 variants. Our results demonstrate that one month of either overexpression of wild-type hTDP-43 or cytoplasmic hTDP-43 expression is sufficient to induce significant cortical and hippocampal gliosis. Although both microglia and astrocytes exhibited morphological signatures of activation, the nature, spatial distribution, magnitude of this response and cellular profile of glial activation are contingent upon both the specific brain region and the subcellular location of hTDP-43.

Our findings challenge a simplistic, binary “resting versus activated” view of microglia, supporting a model of a complex activation spectrum (Butovsky and Weiner, 2018; Masuda et al., 2020; Rohan Walker and Yirmiya, 2016). A recent study in human ALS tissue and the rNLS TDP-43 mouse model reported that microglia display a stage-dependent transition from hypertrophic morphology to a later dystrophic morphology and these changes correlate with increased load of pTDP-43 (Swanson et al., 2025). In the tTA/WT mice described here, PFC microglia exhibited significant soma enlargement, yet Sholl analysis suggested that process complexity might be preserved. This dissociation between soma hypertrophy and process retraction may represent an early or primed activation state, where microglia increase their metabolic activity while preserving their surveillance territory. This phenotype aligns with findings in early-stage models of Alzheimer’s disease and chronic stress, where initial changes in soma size precede dramatic alterations in process morphology (Hinwood et al., 2013; Torres-Platas et al., 2014). In contrast, the more profound process retraction observed in motor and sensory cortices in tTA/WT mice, and globally in tTA/ΔNLS mice, indicates a more advanced shift toward a reactive, potentially phagocytic state. The widespread and severe morphological changes in the tTA/ΔNLS model underscore the potent pro-inflammatory effect of cytoplasmic TDP-43, which is a key pathological hallmark in the majority of ALS/FTD cases (Neumann et al., 2006).

A study on TREM2, a transmembrane protein expressed by microglia, demonstrated its involvement in the phagocytic clearance of pathological TDP-43 (Xie et al., 2022). Moreover, TREM2 deficiency resulted in reduced microglial activation in response to TDP-43 accumulation leading to neuronal damage and motor impairments. The expression levels of TREM2 in the spinal cord and cortex from ALS patients were significantly greater than in control patients. Co-immunoprecipitation essays revealed that TREM2 interacts with hTDP-43 in both mice and humans with ALS (Xie et al., 2022). Recent work from the same group in rNLS8 mice, an ALS model with pan-neuronal cytoplasmic TDP-43 overexpression, showed that TREM2-deficiency decreased the occurrence of rod-shaped microglia, a unique subpopulation involved in modulating excitatory inputs through the engulfment of excitatory synapses, impacting on both neuronal activity and behavioral outputs (Xie et al., 2025).

The early microglial activation in both MC and SSC regions aligns with some features of the clinical presentation of ALS/FTD, where patients can present with both motor and cognitive symptoms (Consonni et al., 2021; Huynh et al., 2020). Our findings suggest that microglial responses might be an early event in disease progression, potentially contributing to the initial phases of neurodegeneration.

Among all glial outcomes assessed, the astroglial response emerged as the most striking discriminator between WT and ΔNLS TDP-43 models. The restricted astrogliosis in tTA/WT mice versus the widespread, robust reaction in tTA/ΔNLS mice suggests that astrocytes are highly sensitive to the presence of cytoplasmic TDP-43. Another mouse model of TDP-43 proteinopathies, prpTDP-43^A315T^, showed that both the loss and gain of function of TDP-43 is sufficient to induce glial reactivity characterized by astrogliosis and microglial activation (Jara et al 2019). This strong astroglial response could have deep consequences in neuronal health (Won et al., 2025). Reactive astrocytes can lose critical homeostatic functions and release pro-inflammatory cytokines, contributing to a toxic environment (Zimmer et al., 2024). Our data indicates that astrogliosis is not just a general marker of neuroinflammation but might be a key indicator of the severity and nature of the underlying proteinopathy. Differences in glial responses found between WT-hTDP-43 and ΔNLS-hTDP-43 models align with recent single-cell transcriptomic studies showing that microglia and astrocytes adopt regionally distinct reactive states in ALS, FTD and AD (Guttenplan et al., 2021; Leng et al., 2021). Such heterogeneity may explain why certain cortical and hippocampal regions are particularly vulnerable in early disease stages.

Glymphatic dysfunction may contribute to ALS pathogenesis, as evidenced by alterations in diffusion tensor imaging studies in patients in comparison to healthy individuals (Baek et al., 2025; Liu et al., 2024). Elevated AQP4 with loss of astrocytic endfeet polarization has been reported in the brainstem and cortex of the SOD1^G93A^ rat model of ALS (Bataveljic et al., 2012). Even though an increase in AQP4 mRNA expression in the spinal cord of SOD1^G93A^ mice was reported as the disease progressed (Dai et al., 2017; Nicaise et al., 2009), AQP4 polarized localization at the endfeet of astrocytes decreased at the disease onset and end stages (Dai et al., 2017).

In hippocampal regions from AD patients, the presence of phosphorylated TDP-43 inclusions was associated with alterations in blood-brain barrier integrity and the reduction in astrocytic AQP4 expression, which correlated with disease severity (Santiago-Balmaseda et al., 2024; Santiago et al., 2025). The reduction in AQP4 area fraction in blood vessels observed in AD patients led these authors to speculate that AQP4 relocates from the astrocytic endfeet to other compartments of the cell, leaving the glymphatic system affected. In agreement with this, another study in TDP-43-depleted astrocytes observed an increase in total AQP4 expression, but mostly dispersed throughout the cytoplasm rather than at the plasma membrane astrocytic endfeet (Khalfallah et al., 2018), pointing out that TDP-43 may modulate AQP4 expression and relocalization.

AQP4 polarization was significantly reduced in hTDP-43-ΔNLS mice, suggesting impaired perivascular astrocytic organization. This alteration could contribute to glymphatic failure, leading to inefficient clearance of misfolded TDP-43 and other neurotoxic aggregates. Consistent with our results showing decreased AQP4 PI in the brains of tTA/ΔNLS mice, disruption of the glymphatic function at early stages of the disease has been described in rNLS8 mice (Zamani et al., 2022). Recent studies by the same group showed that cortical AQP4 mRNA expression was decreased after 7 days of transgene induction (Zamani et al., 2025). Even though altered glymphatic function was detected at 3 days post cytoplasmic TDP-43 induction, its function normalized to control levels by day 7, but cortical AQP4 mRNA expression was significantly altered. These findings highlight early glymphatic dysfunction in ALS, suggesting its potential as a therapeutic target. By 21 days post-induction, glymphatic dysfunction was evident across all brain regions assessed and coincided with widespread degeneration, significant motor impairments and astrogliosis, but AQP4 total expression showed no differences with the control group. Although AQP4 polarization was not assessed in that study, we hypothesize that it might be likely altered, as we observed 4 weeks after CamK2 promoter-driven cytoplasmic TDP-43 overexpression.

The generalized astrogliosis observed in hTDP-43-ΔNLS mice might be both a cause and a consequence of AQP4 dysregulation, reinforcing a cycle of neuroinflammation and glymphatic dysfunction. Differences between models highlight the complexity of TDP-43 pathology, with nuclear vs. cytoplasmic localization shaping the neuroinflammatory and clearance mechanisms.

Although it is unclear whether and how microglial and astrocytic responses might be causally linked, or whether they represent parallel downstream effects of TDP-43 levels/mislocalization, these findings emphasize the importance of glial dysregulation as an early driver of pathology in TDP-43 proteinopathies, with implications for therapeutic strategies aimed at modulating microglial phenotypes. Some limitations of the study include: 1) our data provide insights into region-specific glial activation present at one month after induction; however, temporal relationships to neuronal injury and clearance function remain to be established in future studies. 2) Sex-specific differences in glial activation and glymphatic function have been reported; the current design did not include sex-stratified analyses, limiting generalization to both sexes. 3) Because stainings were performed separately for each line, direct magnitude comparisons between tTA/WT and tTA/ΔNLS were not performed. 4) Regional hTDP-43 neuronal expression and penetrance were not quantified in both lines at this time point; thus, differences in Iba1 and GFAP staining may partly reflect expression load or anatomical targeting, which is not identical in tTA/WT and tTA/ΔNLS lines (Igaz et al., 2011).

Our study reveals that gliosis plays critical but distinct roles in hTDP-43-WT and hTDP-43-ΔNLS mice. While WT-TDP-43 overexpression induces moderate glial activation, cytoplasmic TDP-43 mislocalization leads to widespread gliosis, altered AQP4 distribution, and potential glymphatic impairment. These results highlight the need for therapeutic strategies targeting both neuroinflammation and protein clearance to mitigate TDP-43-related neurodegeneration.

## Acknowledgements

We thank Veronica Risso, Jesica Unger and Lucia Garbini for assistance with animal husbandry, and Agostina Presta and Analía López Díaz for technical assistance.

## Authors’ contributions

Conceptualization: GN, LMI; Data curation: GN, FV, VN, LMI; Formal analysis: GN, FV, LMI; Funding acquisition: LMI; Investigation; GN, FV, AD, VN, LMI; Methodology: GN, FV, VN, LMI; Project administration: LMI; Supervision: AD, LMI; Visualization: GN, FV, LMI; Writing – original draft: GN, FV, LMI; Writing – review and editing: GN, FV, AD; VN, LMI.

## Data Availability Statements

The data underlying this article will be shared on reasonable request to the corresponding author.

## Figure Legends

**Supplementary Figure 1. Subregional analysis of microglial activation in hippocampal CA1 and DG areas from tTA/WT hTDP-43 mice.**

**(A)** Schematic cartoon of hippocampal subregions defined for analyzing microglial features. (B) Representative CA1 and DG images of Iba1+ cells (red) and Hoechst staining (blue) from control and tTA/WT mice. Dotted lines delineate different layers: in CA1, Rad (radiatum layer), Or (oriens layer) and Py (pyramidal cell layer); and in DG, Gr (granular layer) and Po (polymorphic layer). Scale bar: 20 µm. **(C)** Quantification of Iba1+ area in CA1 showed an increase in the layers Or Rad of tTA/WT animals compared to control. **(D)** In the DG, Iba1+ area is increased in both Gr and Po layers. Images were acquired using a Zeiss AxioImager 2 ApoTome 2 system. Statistical analysis was performed using Student’s *t*-test (* p < 0.05, ** p < 0.01). Results are expressed as mean ± SEM. n = 9 - 12 animals per group.

**Supplementary Figure 2. Layer-specific analysis of microglial activation in hippocampal CA1 and DG subregions of tTA/ΔNLS hTDP-43 mice.**

**(A)** Schematic cartoon of hippocampal subregions defined for analyzing microglial features. (B) Representative CA1 and DG images of Iba1+ cells (red) and Hoechst staining (blue) from CA1 and DG in control and tTA/ΔNLS mice. Dotted lines delineate different layers: in CA1, Rad (radiatum layer of the hippocampus), Or (oriens layer of the hippocampus) and Py (pyramidal cell layer); and in DG: Gr (granular layer) and Po (polymorphic layer). Scale bar: 20 µm. (C–D) Quantification of Iba1-positive area normalized to control values reveals a significant increase in CA1 strata and in the granular layer of the DG, whereas no significant differences are observed in the polymorphic layer. Images were acquired using a Zeiss AxioImager 2 ApoTome 2 system. Statistical analysis was performed using Student’s *t*-test (* p < 0.05, ** p < 0.01). Results are expressed as mean ± SEM. n = 9 - 12 animals per group.

