## Supplementary figures and images for "Region-specific features of early glial activation and Aquaporin-4 dysregulation in conditional mouse models of TDP-43 proteinopathies"

### Supplementary Figures 1-2

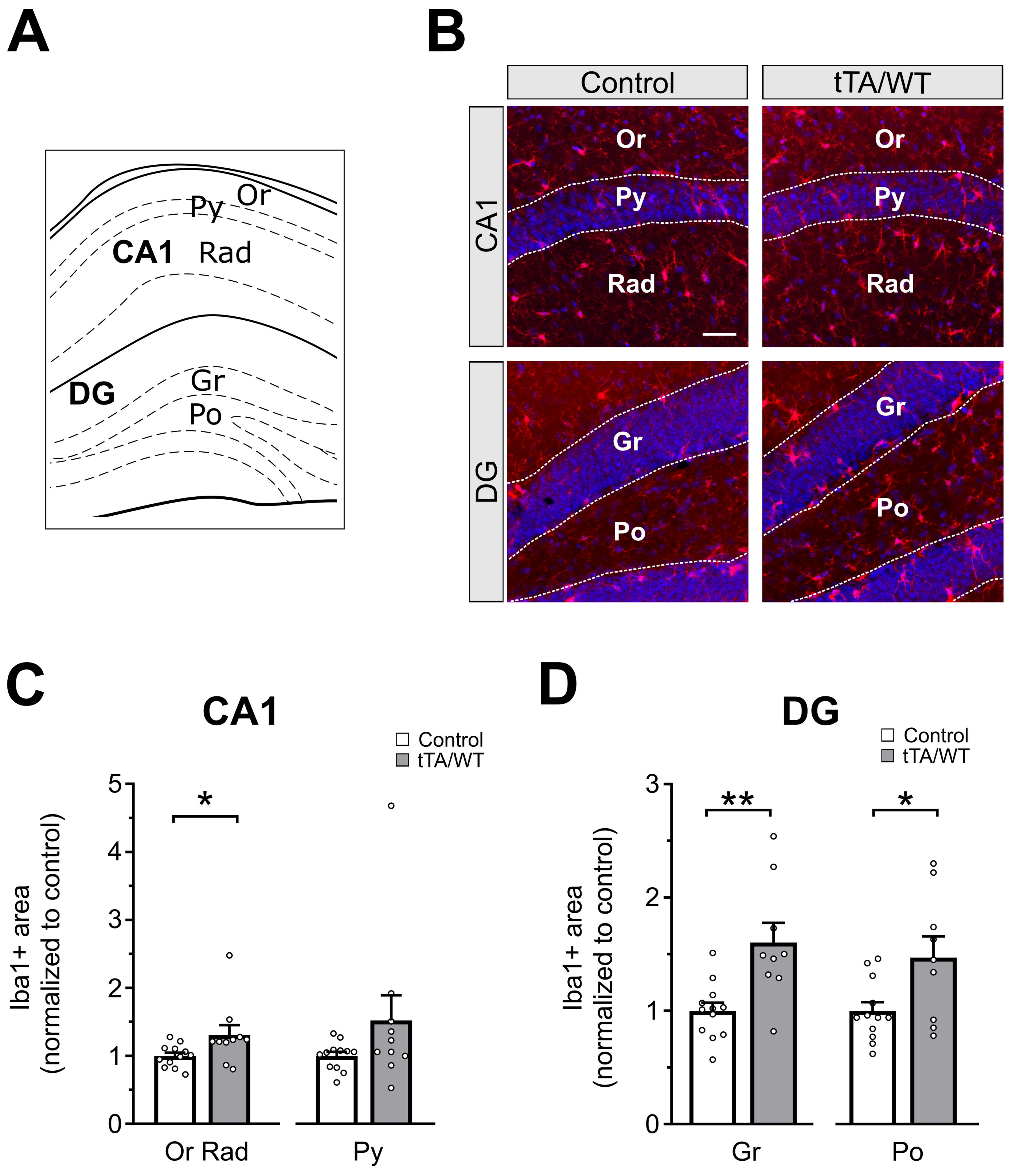

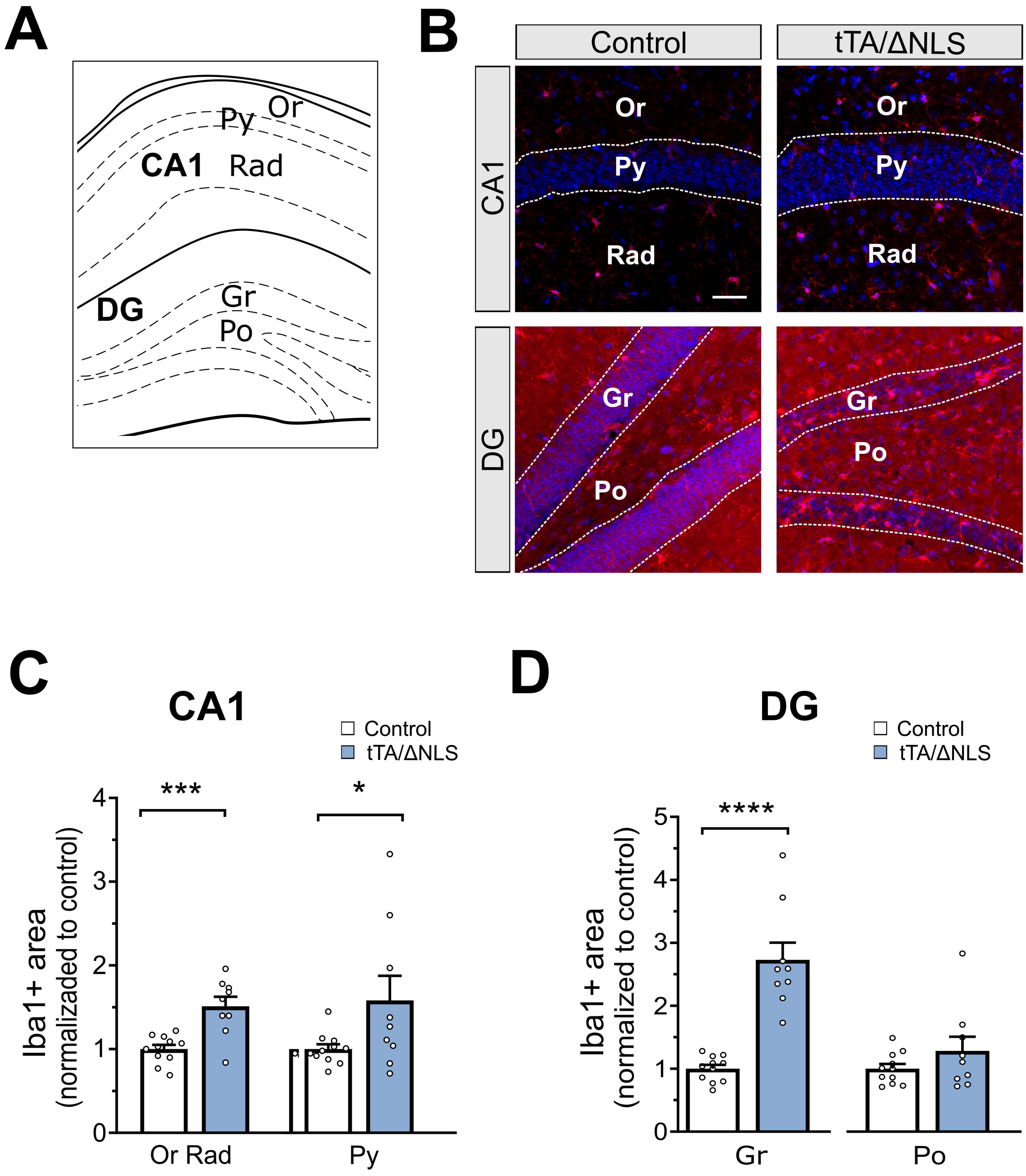
